# Multi-view manifold learning of human brain state trajectories

**DOI:** 10.1101/2022.05.03.490534

**Authors:** Erica L. Busch, Jessie Huang, Andrew Benz, Tom Wallenstein, Guillaume Lajoie, Guy Wolf, Smita Krishnaswamy, Nicholas B. Turk-Browne

## Abstract

The complexity and intelligence of the brain give the illusion that measurements of brain activity will have intractably high dimensionality, rife with collection and biological noise. Nonlinear dimensionality reduction methods like UMAP and t-SNE have proven useful for high-throughput biomedical data. However, they have not been used extensively for brain imaging data such as from functional magnetic resonance imaging (fMRI), a noninvasive, secondary measure of neural activity over time containing redundancy and co-modulation from neural population activity. Here we introduce a nonlinear manifold learning algorithm for timeseries data like fMRI, called temporal potential of heat-diffusion for affinity-based transition embedding (T-PHATE). In addition to recovering a lower intrinsic dimensionality from timeseries data, T-PHATE exploits autocorrelative structure within the data to faithfully denoise dynamic signals and learn activation manifolds. We empirically validate T-PHATE on three human fMRI datasets, showing that T-PHATE significantly improves data visualization, classification, and segmentation of the data relative to several other state-of-the-art dimensionality reduction benchmarks.These notable improvements suggest many potential applications of T-PHATE to other high-dimensional datasets of temporally-diffuse processes.

## 2 Introduction

As we walk through the world, the human brain performs innumerable operations to process, represent, and integrate information. How does a system composed of limited computational units represent so much while handling a constant barrage of incoming information? One possible explanation is that individual neural units operate in tandem to encode and update information in population codes, which exponentially expands the distinct ways information can be represented by a fixed number of units. Neural population codes better predict behavior than single neurons, particularly in how representations change over time in response to new information [1–5]. Recent studies have shown that although neural population codes exist in high ambient dimensions [6]), these dimensions are not orthogonal: they are redundant [7] and from them emerge dominant latent signals that code for behavior and information [8, 9].

Principles of neural population codes were defined using direct neural recordings in nonhuman primates but can be extended to noninvasive, indirect measurements of brain activity using functional magnetic resonance imaging (fMRI). Multivariate pattern analysis [10, 11] of fMRI activation is one such extension that has yielded valuable insights into the structure and content of cognitive representations, including low-level [12] and high-level [11, 13, 14] sensory stimuli, as well as higher-order cognitive processes like memory [15], emotion [16], narrative comprehension [17], and theory of mind [18]. These insights have largely come from group-level analyses, requiring aggregation over multiple subjects and timepoints to draw conclusions because of the high levels of spatio-temporal noise inherent to fMRI. Like other types of high-throughput biomedical data, fMRI noise is pervasive at multiple levels, from subject movement and scanning conditions to measured blood-oxygenation level-dependent (BOLD) signal drift and physiological confounds [19]. fMRI has low temporal resolution relative to the cognitive processes many studies attempt to measure, with typical brain volume acquisition time around 1-2 seconds [20]. Further, the BOLD signal is a slow, indirect measurement of neuronal activity, peaking approximately 4-5 seconds after stimulation before returning to baseline. Finally, complex cognitive processes like memory [21], narrative comprehension [17, 22, 23], or learning are dynamic: they do not exist at a singular timepoint but unfurl over longer periods of time [24]. Such complex yet fundamental processes require integration across modalities and time. The brain likely performs computations on these representations at varying timescales to support conscious experience [25–28].

In sum, many brain representations of interest are coded in high-dimensional patterns of activation [2–4, 10, 11], which can be characterized by low-dimensional latent signals [1, 9, 29]. fMRI affords unique insight into the healthy, behaving human brain, but the data are noisy, sampled slowly, and blurred in time, giving rise to high signal autocorrelation [20, 30]. Further, complex cognitive processes unfurl over time at differing intrinsic neural timescales along the brain’s processing hierarchy [25, 26]. Addressing these issues requires consideration of the various ways in which time and dimensionality interact with the measured BOLD signal with the goal of optimally characterizing fMRI activity at the single-subject level.

Here we introduce “temporal potential of heat-diffusion for affinity-based transition embedding” (T-PHATE) as a nonlinear manifold learning algorithm designed for high-dimensional and temporally-dynamic biological signals. We apply T-PHATE to fMRI data measured during complex cognitive processing, a ripe test-bed as it has known high dimensionality, multi-source noise, and temporal autocorrelation stemming from both measurement sources and the signal of interest. Most previous studies exploring fMRI activity with dimensionality reduction have relied upon linear heuristics to project brain activation patterns into a low-dimensional space [31–33]. Nonlinear dimensionality reduction algorithms have been applied in past studies to characterize the geometry and underlying temporal dynamics in neural recordings [6, 8, 29]) by integrating local similarities among data points into a global representation [9, 34–37], but remain underutilized in fMRI despite the potential to explain age-old questions about cognition[29]. We build upon the manifold learning algorithm PHATE [34] designed for high-throughput, high-noise biomedical data, with a second, explicit model of the temporal properties of BOLD activity. This second view is learned from the data to capture the temporal auto-correlation of the BOLD signal in different brain regions and the dynamics specific to a given stimulus (Fig. 1)A).

**Figure 1:**
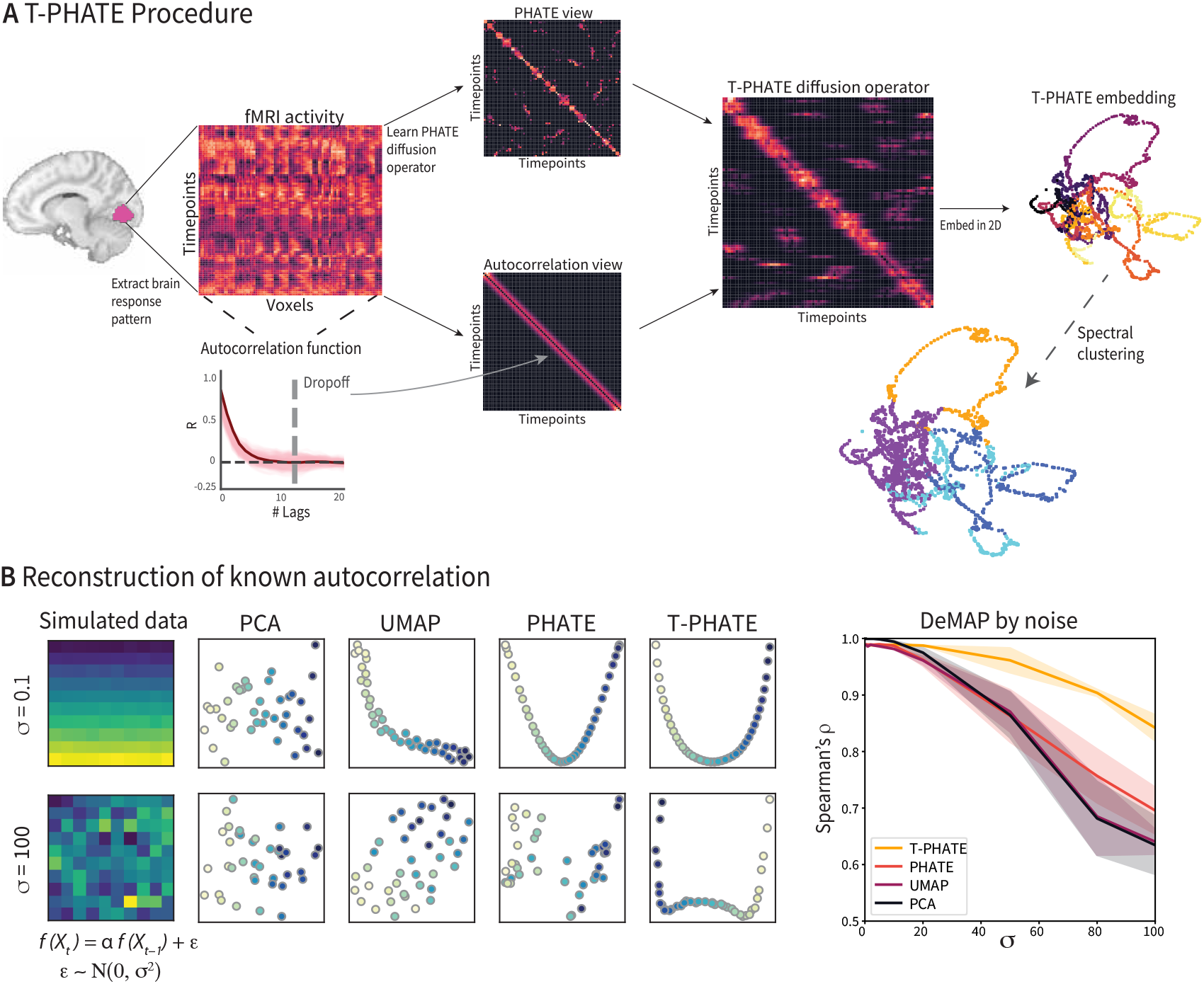
**A**. Temporal PHATE (T-PHATE) procedure. A multivariate pattern of fMRI activation over time (timepoints by voxels) is extracted from a region of interest (ROI). PHATE is then run to learn a PHATE-based affinity matrix between timepoints. The autocorrelation function is also estimated for the data separately to make an autocorrelation-based affinity matrix. These two views are combined into the T-PHATE diffusion operator then embedded into lower dimensions. **B**. We tested this approach on simulated data with a known autocorrelative function and varied the error added to the signal. T-PHATE both visually better reconstructs the autocorrelative signal from the noise and quantitatively better preserves manifold distances, compared with PCA, UMAP, and PHATE. See Figure S1A for additional benchmarks.

Using a manifold preservation metric called DeMAP [34], we benchmark T-PHATE against principal components analysis (PCA) and uniform manifold approximation (UMAP) [38], dimensionality reduction methods that have been commonly applied to fMRI data [31, 32, 39, 40], as well as PHATE [34], a reduced form of T-PHATE excluding the second, temporal view. We compare T-PHATE with locally linear embeddings (LLE; [41, 42]), isometric mapping (Isomap; [35]), and the popular visualization method t-Distributed Stochastic Neighbor Embedding (t-SNE; [43]) as additional benchmarks in supplementary material as noted in each main figure caption. Unlike prior studies [29]), our goal is to learn a manifold that captures temporal dynamics within a naturalistic task that could later be extended to new subjects [44] or used to predict future brain states. Thus, we test T-PHATE on two movie-viewing fMRI datasets and find that it both denoises the data and affords enhanced access to brain-state trajectories than either voxel resolution data or a variety of other dimensionality reduction approaches. By mitigating noise and voxel covariance, this subspace yields clearer access to regional dynamics of brain signals, which we can then relate to time-dependent psychological phenomena. In all, T-PHATE reveals that information about cortical dynamics during naturalistic stimuli lies in a low-dimensional latent space best modeled in nonlinear, temporal dimensions.

## 3 Results

### 3.1 Evaluating manifold quality

fMRI is a powerful tool for studying the neural bases of higher-order cognitive abilities. fMRI data are highly noisy in both space and time, with the measured BOLD signal canonically peaking 4-5 s after stimulus onset before slowly returning to baseline. Naturalistic, complex stimuli like movies are increasingly used to probe real-world cognition and reasoning in the brain. These stimuli are dynamic, with conversations and plot lines playing out over different timescales, which cause the autocorrelation of brain responses to extend beyond the curve of the BOLD signal and vary according to the temporal integrative properties of individual brain regions. Thus, we designed T-PHATE as a variant of the PHATE algorithm that combines the robust manifold-geometry preserving properties of PHATE with an explicit model of the data’s autocorrelation in a dual-diffusion operator. We compared manifolds learned with T-PHATE to those from PCA, PHATE, and UMAP. We validated the quality of manifolds learned from data with a simulated high auto-correlative signal using Denoised Manifold Affinity Preservation (DeMAP)[34]. DeMAP takes pristine, noiseless data (here, a simulated multivariate timeseries where *f* (*X*_*t*_) = *α f* (*X*_*t*_ − _1_)) and computes the Spearman correlation between the geodesic distances of the noiseless data and the Euclidean distances on the embeddings learned from noisy data. Noisy data were generated by adding an error term *ϵ* to the pristine data, where *ϵ*∼ *N* (0, *σ*^2^). We tested robustness to noise by varying *σ*= (0, 100). Higher DeMAP scores suggest that an embedding method performs effective denoising and preserves the geometric relationships in the original data. DeMAP results for PCA, UMAP, PHATE,and T-PHATE are shown in Fig. 1B, and in Fig. S1A for LLE, Isomap, and t-SNE. At low levels of noise, all methods achieve high DeMAP scores, though PCA visualizations do not appear to reflect much structure. With increasing noise, T-PHATE outperforms the other methods at denoising the simulated data, providing visualizations that most closely resemble that of lower noise conditions.

Having validated that T-PHATE can learn meaningful manifolds from simulated data even under noisy conditions, we next applied T-PHATE to fMRI data. First, we embedded the movie-viewing data from the *Sherlock* and *StudyForrest* datasets with PCA, UMAP, PHATE, and T-PHATE in two dimensions to visually inspect the properties of the data highlighted by the manifold. Visually, T-PHATE embeddings denoise the timeseries data and capture stimulus-related trajectory structure better than the other methods, as shown in all brain regions for a sample subject (Fig. 2A) as colored by sequential timepoint. PCA visualizations show no apparent structure or clustering, whereas UMAP shows at least slight clustering of nearby timepoints in high visual (HV) and early auditory (EA) cortices but often creates small, shattered clusters with local structure placed through the latent space. PHATE visually yields slight improvements over UMAP, notably in HV and EA. Clusters in these plots appear larger and less disjointed, connected by tendrils that flow from a hub-like center space where the majority of clusters exist in the embedding. T-PHATE, however, reveals remarkable trajectories through the latent space that are clearly reflective of temporal structure but also shows a hub in the center of the space. T-PHATE manifolds also better reflect non-temporal attributes of the movie, as shown by classification of stimulus features at given timepoints (Fig. 2B). We ran a support vector classification to predict whether music was playing at a given timepoint from the embeddings of neural data (and as a baseline, the original voxel-resolution data). Significance of these predictions was tested by shifting the labels in time with respect to the brain data. Results are presented as the z-statistic of the true prediction accuracy between brain data and movie labels normalized to the mean and standard deviation of the null distribution from the shifted versions.

**Figure 2:**
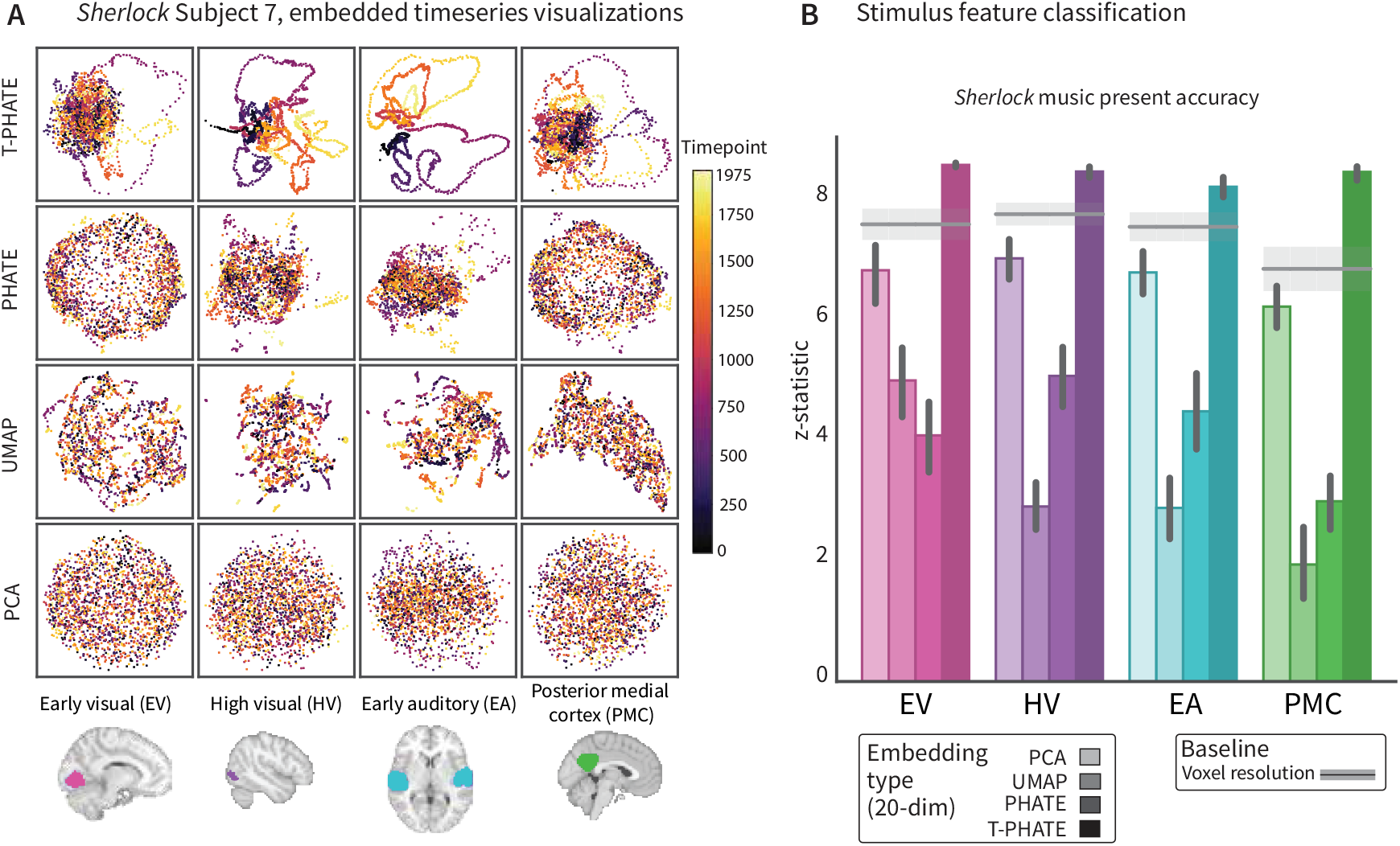
**A**. Manifold embeddings using PCA, UMAP, PHATE, and T-PHATE for the *Sherlock* dataset in an example subject (7) and 4 regions of interest. Points are colored by time index. **B**. Support vector classification accuracy predicting the presence of music in the show at the corresponding timepoint. Results are presented as a z-statistic of the true accuracy versus a null distribution of accuracies from time-shifting the timeseries (which preserves their autocorrelation). Extended SVC results included in Supplemental Fig. 2

The clarity of the dynamic structure recovered by T-PHATE prompted two follow-up questions. First, does the performance of T-PHATE depend on the use of autocorrelation to represent temporal structure during manifold learning? To answer this question, we tested two alternatives to T-PHATE, PHATE+Time and Smooth PHATE. PHATE+Time adds the TR label for each timepoint as an additional feature vector to the voxel data before embedding with PHATE. PHATE+Time manifolds show the clearest temporal trajectory through the embedding space (Fig. 3A), unraveling the brain data in time due to the disproportionate weighting of time in the input data, with less of the hub struc-ture shown by T-PHATE. Smooth PHATE performs temporal smoothing along each voxel with a double gamma function (the canonical hemodynamic response function), which is a common preprocessing step for fMRI data generally, before embedding with PHATE. Smooth PHATE yields comparable structure to PHATE or UMAP, with shattered trajectories and mild clustering.

**Figure 3:**
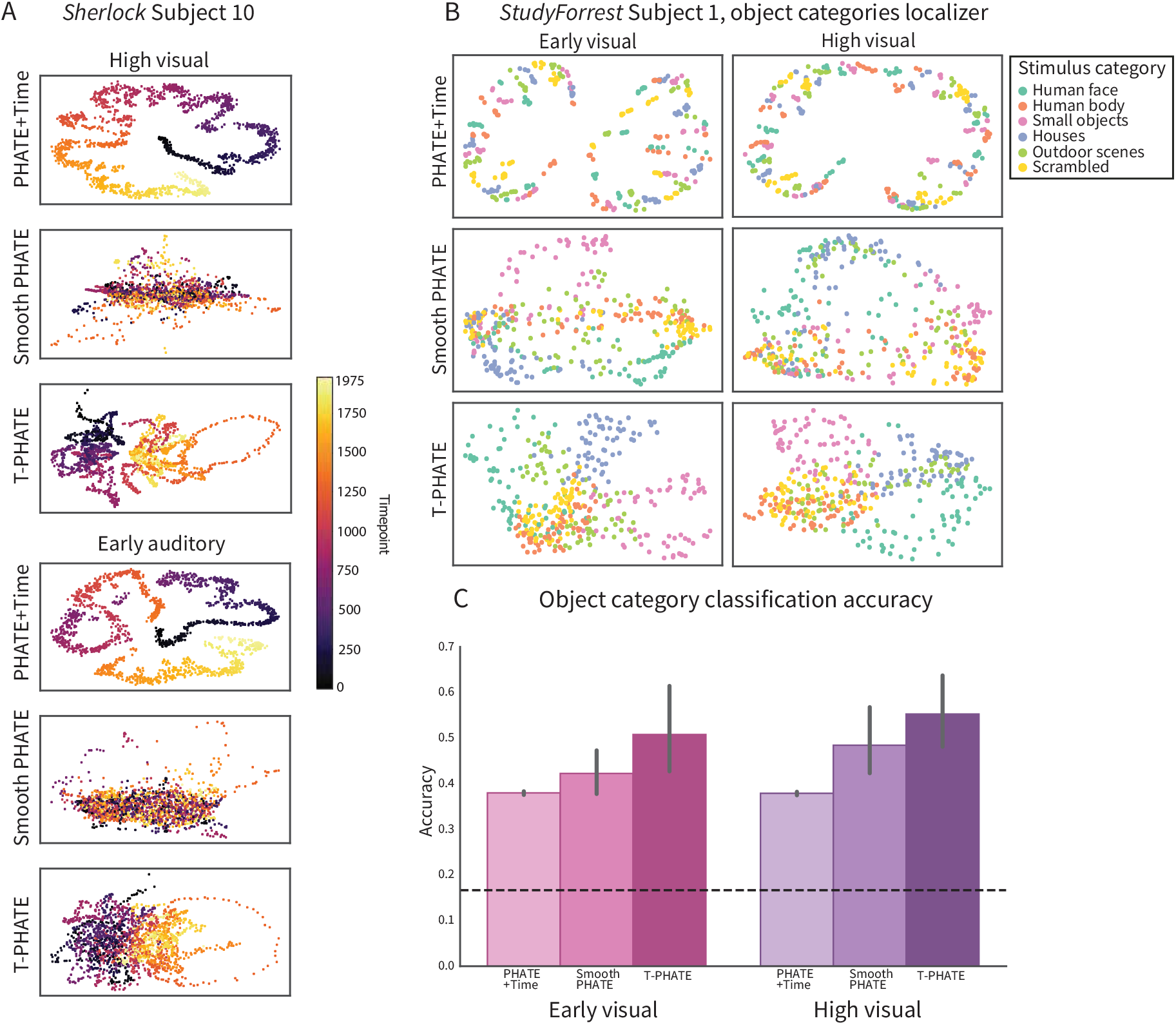
**A**. Manifold embedding of *Sherlock* data in high visual (HV) and early auditory (EA) ROIs with T-PHATE, PHATE+Time, and Smooth PHATE, colored by time. **B**. *StudyForrest* localizer data embeddings using T-PHATE, PHATE+Time, and Smooth PHATE in early visual (EV) and high visual (HV). T-PHATE reveals the best clustering by stimulus category, showing that the manifold successfully denoises data to learn a more accurate manifold. **C**. T-PHATE manifolds also afford better stimulus category classification accuracy. Bars represent average accuracy across subjects; error bars represent 95% CIs of the mean from bootstrap resampling; dashed line represents chance. See Fig. S1B for additional benchmark results.

Second, are the benefits of T-PHATE for stimulus feature classification specific to fMRI tasks that are intrinsically linked to time, such as movie-watching? To answer this question, we tested T-PHATE on an object category localizer task from *StudyForrest* with no temporal structure, during which static images of faces, bodies, small objects, houses, scenes, and scrambled images were presented in a random order. We embedded these data with T-PHATE, PHATE+Time, and Smooth PHATE for early visual (EV) and high visual (HV) ROIs. Because of the lack of temporal structure, a meaningful manifold of fMRI activity in visual regions during this task would show a time-independent clustering of categories. T-PHATE manifolds show the best clustering of stimulus categories in the embedding space (Fig. 3B), compared with PHATE+Time and Smooth PHATE. Smooth PHATE showed some clustering but less pronounced, whereas PHATE+Time showed minimal clustering by category but instead still retained temporal structure. We quantified the ability to distinguish object categories in the latent space using a within-subject support vector classifier with leave-one-run-out cross-validation (6-way classification; chance = 1/6) (Fig. 3C). In EV, classification accuracy on T-PHATE embeddings surpassed all other dimensionality reduction methods (T-PHATE M=0.51, CI=[0.42, 0.6]; voxel M=0.46, CI=[0.38, 0.55]; PCA M=0.43, CI=[0.34, 0.52]; UMAP M=0.4, CI=[0.39, 0.42]; PHATE M=0.39, CI=[0.37, 0.42]; PHATE+Time M=0.38, CI=[0.37, 0.39]; Smooth PHATEM=0.42, CI=[0.38, 0.47]). In HV, T-PHATE again outperformed the other dimensionality reduction methods (T-PHATE M=0.55, CI=[0.47, 0.62]; voxel M=0.53, CI=[0.46, 0.61]; PCA M=0.48, CI=[0.46, 0.51]; UMAP M=0.42, CI=[0.41, 0.44]; PHATE M=0.43, CI=[0.42, 0.45]; PHATE+Time M=0.38, CI=[0.37, 0.39]; Smooth PHATE M=0.48, CI=[0.41, 0.56]; additional benchmarks shown in Fig. S1B.). This analysis showed that T-PHATE captures structure in brain activity via autocorrelative denoising, whereas adding time explicitly into the manifold does little for capturing the data geometry aside from unrolling the signal in time. Temporal smoothing provides an advantage over time-agnostic dimensionality reduction (PCA, PHATE, UMAP), but less of an advantage than the autocorrelation kernel in both the dynamic movie task and the non-temporal localizer task.

### 3.2 Event segmentation

We designed T-PHATE to incorporate temporal dynamics (namely autocorrelation) into a manifold learning algorithm for two reasons. First, as shown above, this helps denoise fMRI data given its spatiotemporal noise and improve subsequent analyses such as feature classification. Second, this may help recover cognitive signals that are represented over time in fMRI data. Many cognitive processes operate on longer timescales, including our ability to segment experience into events [45–47]. That is, we do not perceive the world as transient from moment to moment, nor as amorphous and fully undifferentiated, but rather as a series of coherent and often hierarchically structured periods of time [48, 49]. For example, consider the steps involved in taking a flight — traveling to the airport, passing through security, boarding, flying, deplaning, an transiting to the destination. These mental events are reflected in stable states of the brain, which can be captured with a Hidden Markov Model (HMM) to identify the boundaries between events [50–52]. We hypothesized that an HMM would be better able to perform this event segmentation after embedding fMRI data into a manifold learned with T-PHATE. This would suggest that T-PHATE can increase sensitivity to the neural dynamics associated with cognitive processing of the stimulus.

We started by learning the optimal *number* of events experienced by each ROI using a leave-one-subject-out cross-validation procedure on the voxel resolution data (Fig. 4A). We find a pattern of results consistent across datasets where the number of events *K* represented by a brain region is higher for early sensory cortices (EV) than late sensory cortices (HV) and integrative cortices (PMC) (values presented as mean number of events ± s.d. across subjects): *Sherlock*: EV = 53 ±12, HV = 21± 7, AE = 18 ±5, PMC= 22± 6; *StudyForrest*: EV = 111 ±43, HV = 69± 17, AE = 74± 18, PMC = 96 ±24. After fixing the *K* parameter for each ROI based on the voxel data, we learned the number of manifold dimensions that would best reveal temporal structure for each region, subject, and method with the same cross-validation procedure. Number of dimensions is low (between 3-6 dimensions) with small variance across subjects and method. This indicates substantial redundancy, covariance, and noise among voxels in the ROIs, obscuring meaningful, dominant signals (Supplemental Fig. 1).

**Figure 4:**
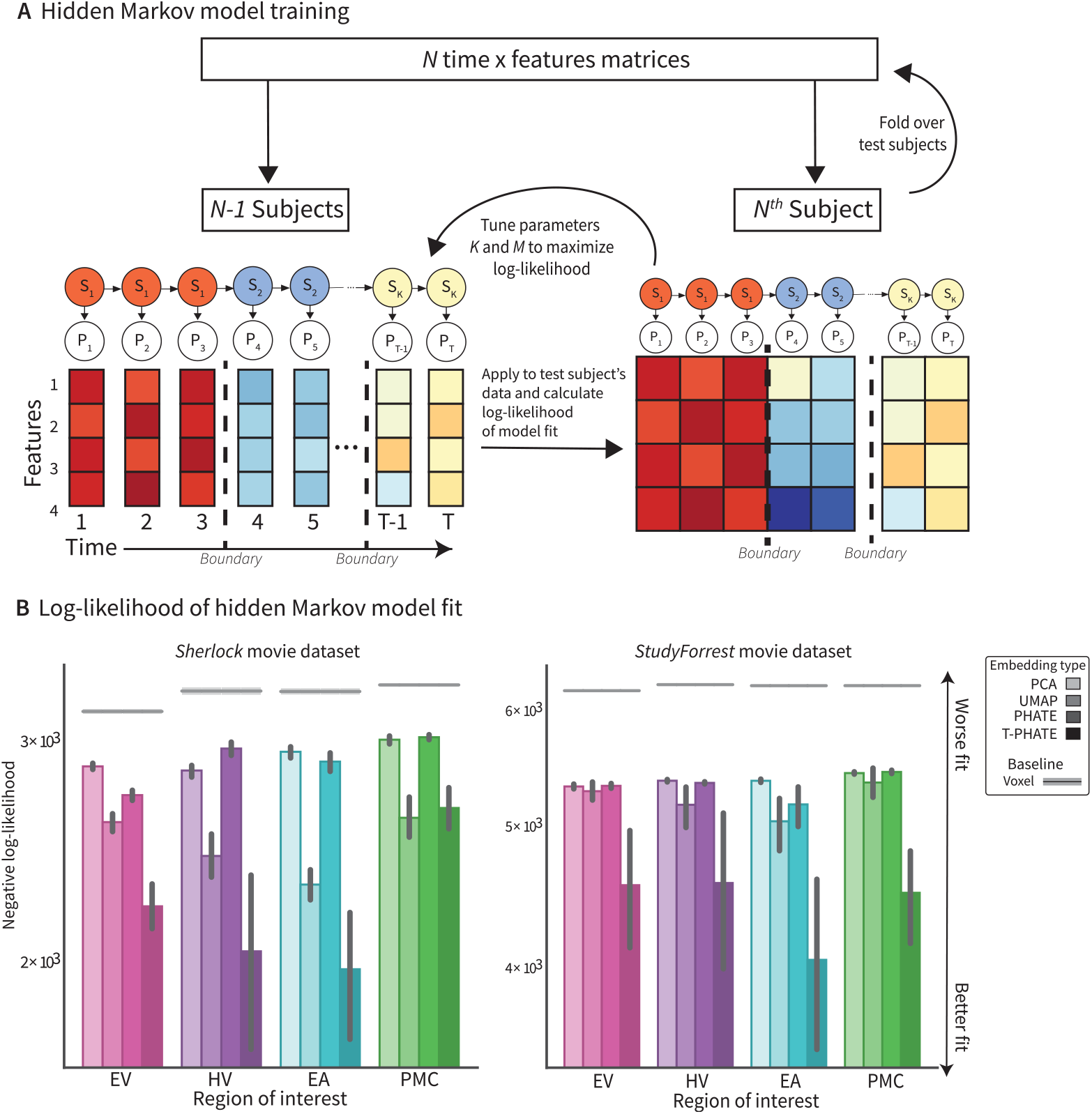
**A**. The HMM is trained on the average data for *N*− 1 subjects to find *K* stable patterns of voxel activation, where the voxel patterns *P* for each timepoint belong to a given hidden state *S*. Model fit is evaluated on a test subject’s data then repeated with leave-one-subject-out cross-validation. Event boundaries (dashed lines) are identified by the model as timepoints at a transition between hidden states. **B**. Log-likelihood of model fit is tested on a given subject using a HMM with hyper-parameters *K* and *M* tuned via leave-one-subject-out cross validation. Model fit here is presented as the negative log-likelihood, where a better model fit results in a *lower* negative log-likelihood. Different dimensionality-reduced approaches are shown as bars and the voxel resolution benchmark is shown as line with shading representing mean and 95% confidence interval.

A goal of T-PHATE is to recover dynamic cognitive signals that may have been previously obscured by the noise of redundancy and noise of the data. If dynamical structure during movie watching is unveiled in the T-PHATE latent space, we would see better success of the HMM on data embedded with T-PHATE than to the other methods or the voxel resolution data. This can be framed as: (1) how well does the model fit the data and (2) how well do boundaries learned by the model capture structure of the neural data. For each subject, ROI, and embedding method, we fit a HMM with *K* hidden states on a *M*-dimensional embedding (where hyper-parameters *K* and *M* were cross-validated at the subject level for ROIs and methods, respectively). Model fit is quantified after hyper-parameter tuning using log-likelihood. T-PHATE shows significantly better model fit across both the *Sherlock* and *StudyForrest* datasets in almost all regions (except PMC for *Sherlock*), visualized as negative log-likelihood, where closer to zero means a better model fit (Fig. 4B). This indicates a higher probability of the HMM’s learned segmentation fitting the test data. To validate whether the boundaries identified actual differences in the data, we then applied the boundaries learned during the HMM fitting procedure to the data and asked whether the neural data from pairs of timepoints that fall within the same event are more similar than pairs that fall across different events (equating temporal distance) (Fig. 5A). T-PHATE embeddings result in significantly better within-versus-between event boundary distinctions in almost all regions (except EV for *StudyForrest*, where T-PHATE is marginally greater than PCA) (Fig. 5B). In the *Sherlock* dataset (and 2/4 ROIs of the *StudyForrest* data), T-PHATE at least doubles the performance of the second-best embedding type. Notably, all dimensionality reduction outperform the original voxel resolution data. This indicates that, though there is an inherent advantage in revealing dominant signal structure in data in lower dimensions, the temporal diffusion step of T-PHATE captures dynamic signals beyond a temporally-agnostic manifold.

**Figure 5:**
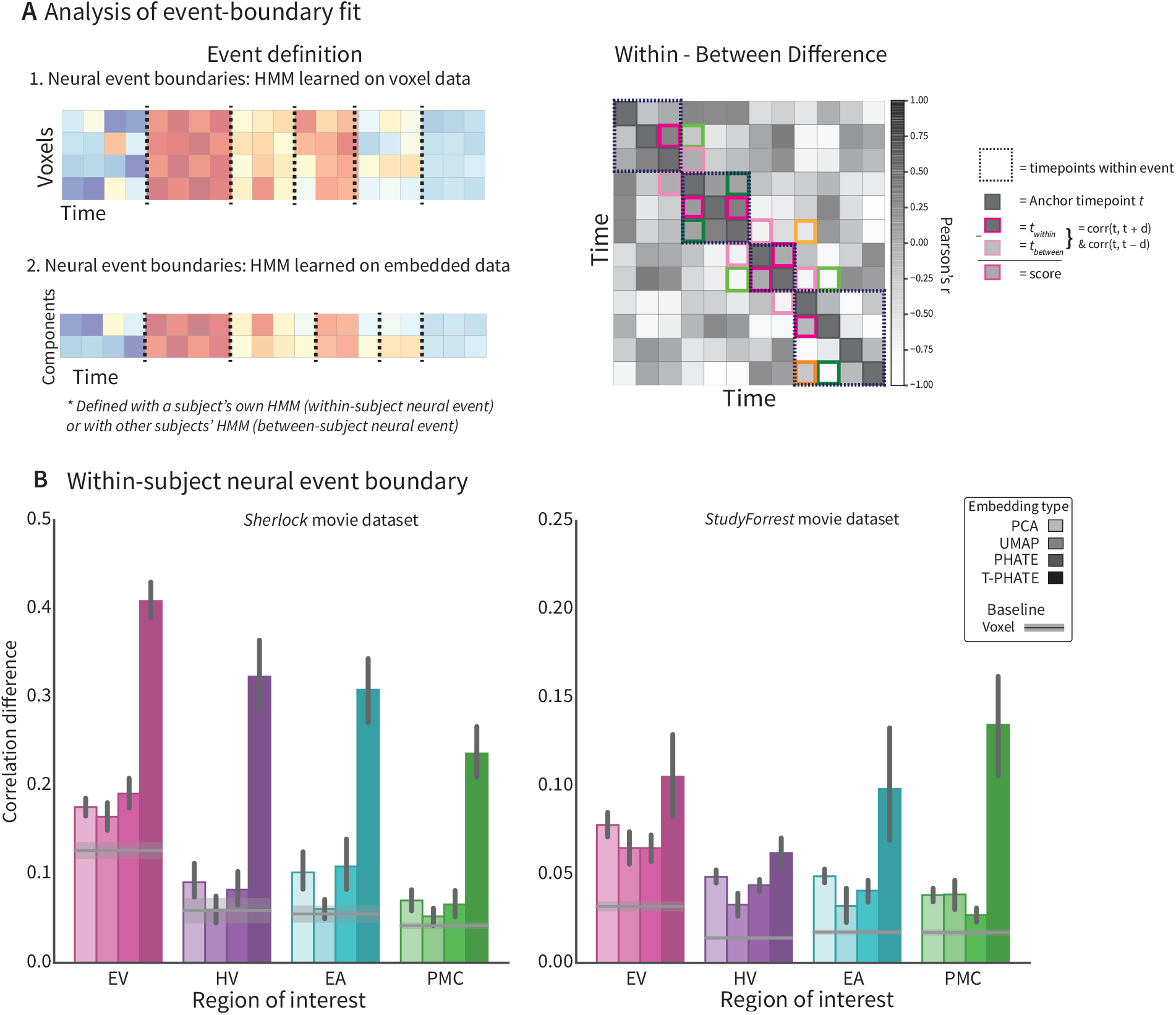
**A**. Event boundaries are learned both from the data (by applying the HMM on voxel resolution or embedded timeseries data and applying those boundaries either within or across subjects) and from independent sources including human behavioral ratings. (Right) Representation of dynamic information in a latent space is quantified by applying event boundaries to embedded data. Data emphasizes brain state transitions if the boundaries maximize the difference between brain activity at timepoints across events versus within events. For each TR *t*_0_ in the timeseries, we correlate the activation pattern across features at *t*_0_ with that of *t*_1_ and *t*_2_ such that *t*_1_ is in the same event as *t*_0_ (within a dashed box) and *t*_2_ is in a different event. The within-between score is the subtraction of *corr*(*t*_2_, *t*_0_) from *corr*(*t*_0_, *t*_1_). To account for intrinsically higher correlations of proximal timepoints, we restrict comparisons to where| *t*_0_ −*t*_1_ = *t*_0_ −*t*_2_|. Here, *corr*(*t*_0_, *t*_1_) is denoted by a darker colored box and its corresponding *corr*(*t*_2_, *t*_0_) a lighter shade. **B**. Neural event boundaries for subject *N* are identified with the HMM and then applied to calculatethe subject’s within-between score. Different dimensionality-reduced approaches are shown as bars and the voxel resolution benchmark is shown as line with shading representing mean and 95% confidence interval. See Fig. S5A for additional benchmark results for this analysis.

In the previous analysis, event boundaries are learned and applied from a HMM fitted within-subject. To ensure the high performance in this analysis was not driven by overfitting from training/testing the boundaries in the same subject, we performed cross-validation between subjects by applying the event boundaries identified for *N*− 1 subjects to subject *N*’s data individually. In other words, for each of *N* subjects, we ran the analysis *N*− 1 times using each of the other subject’s boundaries. Again, we found the strongest effect for T-PHATE relative to the other embedding types (Fig. 6), albeit weaker than when the event boundaries were learned within-subject, as expected. Interestingly,the voxel resolution data outperformed each of the methods other than T-PHATE in a few cases (voxel *>* others in: *Sherlock*: HV UMAP and PHATE, EA UMAP, PMC PHATE; *StudyForrest*: EV PHATE, HV PCA and PHATE, PMC PCA, PHATE and UMAP). This was not seen in the within-subject version, suggesting that some overfitting to subject-specific latent spaces occurred for the other dimensionality reduction methods.

**Figure 6:**
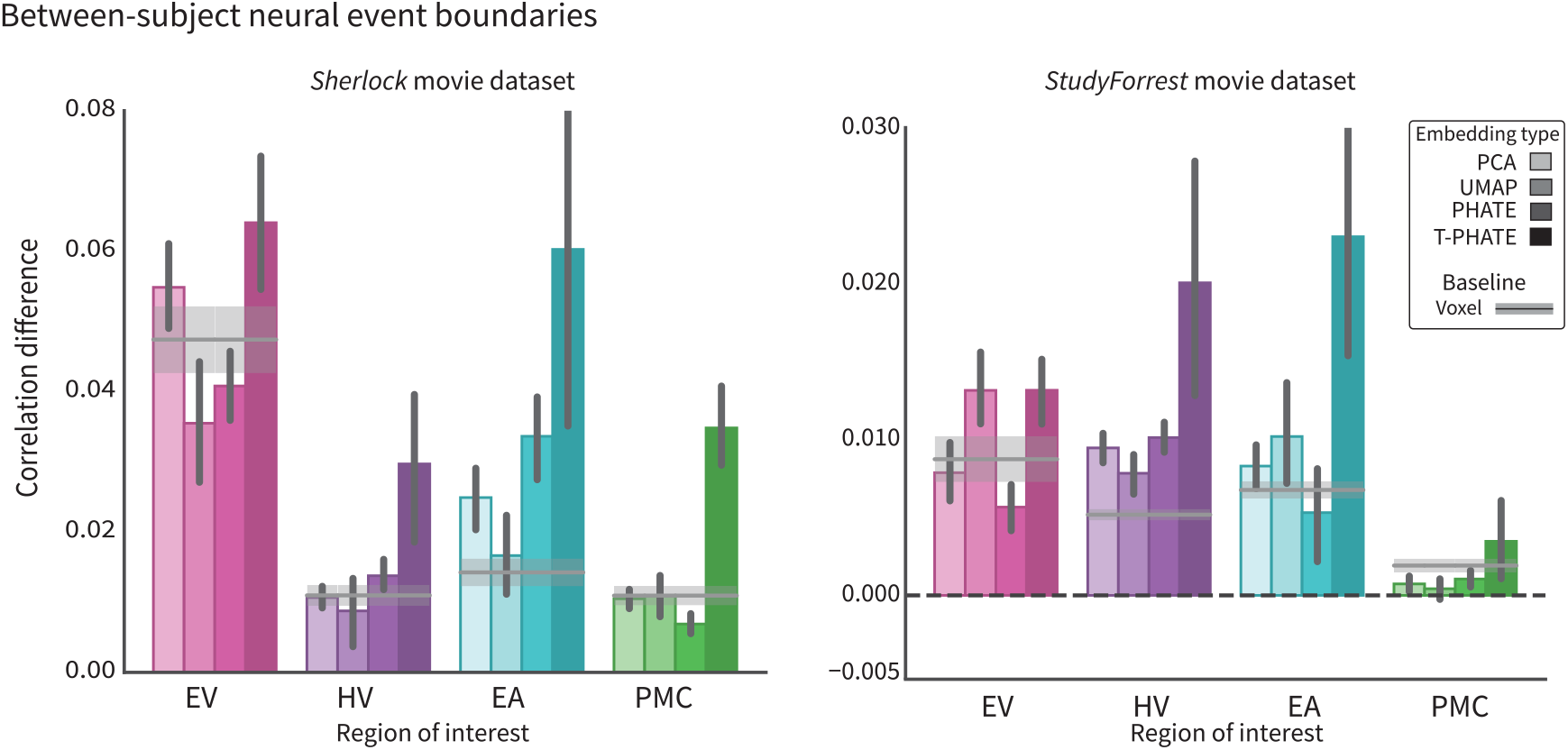
Event boundaries were rotated between subjects by testing subject *N* iteratively with the event boundaries learned from HMMs applied to each of the remaining *N-1* subjects (within ROI and embedding type). T-PHATE embeddings reveal more robust event-related structure even when event boundaries are learned from different subjects. See Fig. S5B for additional benchmark results for this analysis.

Identifying neural event boundaries with HMM is a data-driven approach to understanding how the brain makes sense of continuous stimulation. However, the identified boundaries have no ground truth. Another way to assess whether learned neural boundaries are meaningful is to use a separate behavioral task where individuals watch the same movie and indicate where they believe something changed and there was a shift to a new event. Ample literature supports the use of this type of parsing task for studying human event segmentation [45–47, 53], allowing us to treat human-rated boundaries as a ground truth. A separate cohort of human subjects watched the *Sherlock* movie and were asked to indicate where they thought an event boundary occurred (Fig. 7A). We applied these behavioral boundaries to the neural data in the same fashion as we did the neural events and measured the neural distance within and across boundaries (again equating temporal distance). The ground-truth behavioral event boundaries resulted in significantly stronger within-versus-between neural differences when applied to the T-PHATE embedding than to any of the other methods (Fig. 7B). The PMC is a higher-order integrative region that has previously been shown to predict behavioral boundaries [50, 54]. We replicated these findings, showing a significant effect in PMC on voxel data (and with other methods), but critically T-PHATE improved on this performance by an order of magnitude. This gain is particularly striking because the behavioral data being predicted were generated by other subjects, in a different task, and did not rely on an HMM of neural data to find the boundaries.

**Figure 7:**
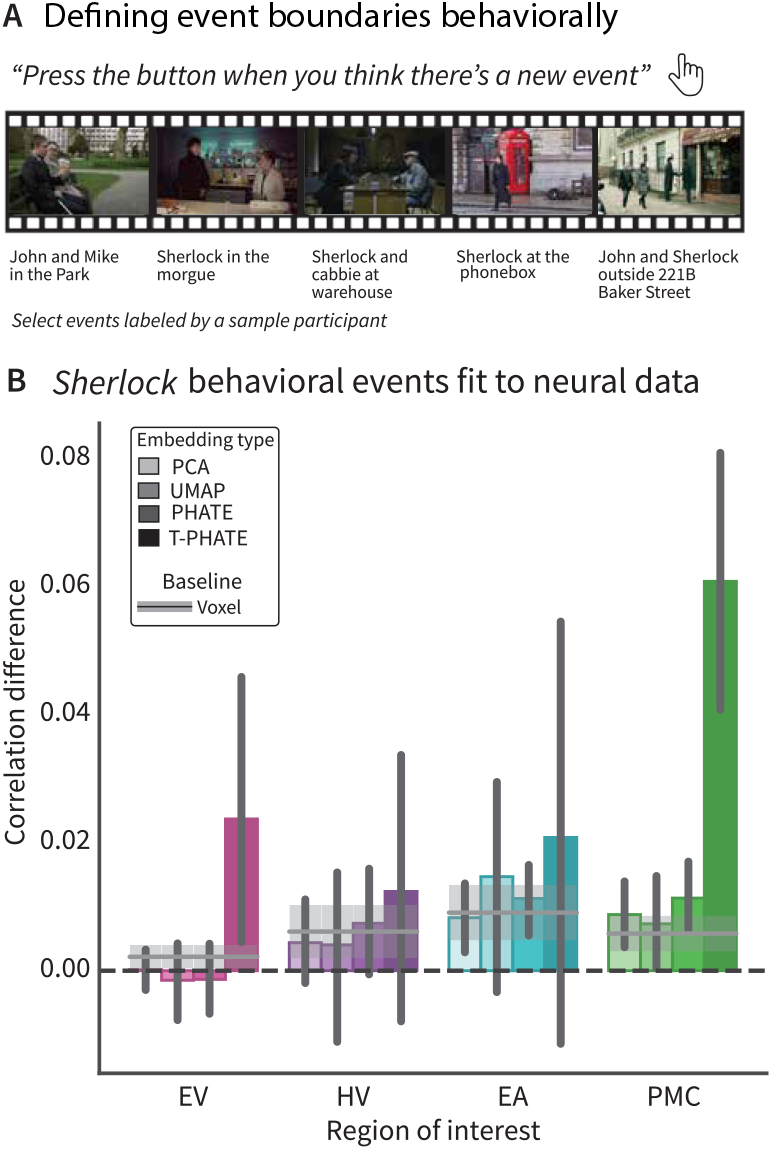
**A**. Event boundaries were also identified by behavioral ratings in a separate experiment on the *Sherlock* dataset with other subjects. Behaviorally-defined event boundaries were then applied to the neural data and within-versus-between event correlation was evaluated as in Fig. 5A) **B**. These behavioral boundaries best mirror event structure in PMC and are best revealed with T-PHATE. Different dimensionality-reduced approaches are shown as bars and the voxel resolution benchmark is shown as line with shading representing mean and 95% confidence interval.

## 4 Discussion

In this study, we investigated the latent space of regional brain activity during naturalistic viewing from fMRI data. We found that this latent space best captured dynamic information about the experience of a stimulus using a low-dimensional subspace identified with T-PHATE, our novel method that incorporates the temporal dynamics of a signal to learn a more robust manifold. By reducing into nonlinear dimensions that capture not just the geometry of multivoxel activity patterns but also their temporal dynamics, we reveal trajectories through cognitive states that were previously obscured by considering the data in its full resolution. T-PHATE can capture meaningful cognitive trajectories in as few as three dimensions from single-subject fMRI data, which is inherently noisy in both space and time.

We quantified the degree to which manifold representations of brain activity reflect the dynamics of the stimulus and cognitive processing using a neural event segmentation framework. We applied Hidden Markov Models to learn transitions between stable neural states that reflect boundaries between event representations in different brain regions. The event boundaries learned on data embedded with T-PHATE provided a better fit to the neural data of same subject, predicted the neural similarity of timepoints from that subject, and generalized to other subjects. Moreover, the data embedded with T-PHATE were able to predict where in the movie other human raters would indicate behaviorally that they sensed an event boundary subjectively. To our knowledge, previous investigations of neural event segmentation have not used dimensionality reduction to either denoise neural data or model the structure of neural dynamics. All dimensionality reduction methods we presented did a better job at modeling event dynamics than the voxel-dimensional data (Fig. 4B), likely because they denoised the data in low dimensions. Nevertheless, T-PHATE vastly outperforms PHATE, UMAP, and PCA. T-PHATE manifolds reveal neural dynamics previously obscured by the high-dimensionality, noise, and redundancy of both fMRI data and the neural population activity underlying it by modeling the latent spatiotemporal geometry of brain activity.

Importantly, the advantage T-PHATE gains in dynamic modeling is not at the expense of static decoding. Multivariate pattern classification of stimulus features on low-dimensional T-PHATE embeddings outperform or match decoding from the voxel resolution data (Fig. 2B; Supplemental Fig. 2) and outperform the other dimensionality reduction methods we consider. This shows that while both time-considerate and time-agnostic manifolds can encode static information, only the former can additionally unveil dynamic information.

Cognitive processes like learning, language comprehension [22], theory of mind [55, 56], narrative understanding [17, 22, 23], and memory [21]. do not occur at a single point in time — they require the accumulation of information over longer temporal windows [24]. These automatic processes define our conscious experience of the world in real-time, structuring continuous input into meaningful events that shape perception and memory [45–47, 50, 57, 58]. To understand how such complex, integrative processing can happen so seamlessly during conscious experience, it is helpful to consider the geometry of event representations in the brain. The fMRI literature suggests that flexible population codes underlie complex cognitive processing [13, 18, 50, 54, 59]. Further, the animal literature shows that population representations underlying cognitive processes can be modeled in low-dimensional manifolds [3, 8, 9], and the temporal unfoldings of behavior are best predicted with nonlinear, temporally-dependent manifolds [8, 60].

To date, linear manifolds have been used to track task-related brain states across block-design cognitive tasks [31, 32] and nonlinear manifolds have been used to model the dynamics of cognition across these tasks [29]. These approaches have required aggregation of whole-brain data across dozens of subjects to define a manifold, and the dynamics captured are coarse, population-average representations of traditional, task-defined cognitive states (such as working memory or social tasks [29], limiting applicability in understanding fine-scale, individual brain dynamics. To our knowledge, no one has yet investigated individual subject manifolds learned from single-subject fMRI data collected with naturalistic stimuli to investigate temporal dynamics of individual brain regions. This is likely due to the noise of single-subject, naturalistic fMRI data. With the proper choice of dimensionality reduction algorithm [6, 9], we accounted for these challenges to unveil latent dynamics of the cognitive processes related to movie watching.

In conclusion, we found that nonlinear, temporal manifold learning can identify a low-dimensional subspace of brain activity from continuous, naturalistic stimuli that captures higher-order cognitive representations in the brain. Critically, latent dimensions learned with this combination of nonlinear dimensionality reduction and temporal diffusion out-performs methods lacking either of those components. This suggests that the relation between the dimensions of individual voxels and the dominant signals that emerge from them are tied by nonlinear relations. By modeling a subject’s BOLD timeseries in these dimensions, we can gauge the trajectory of their brain states throughout the course of the experiment, and we can relate this trajectory to theoretical frameworks like event segmentation theory that facilitate higher-order cognitive processes in the brain. Because this is performed at the level of individual subjects, future investigations could use this approach to probe individual differences, developmental trajectories, and/or clinical disorders in the native latent space. The characteristics of these manifolds could inform theories about the nature and dynamics of human cognition.

## 5 Methods

### 5.1 Manifold embeddings

Different dimensionality reduction algorithms focus on different aspects of a dataset, thus amplify distinct signals depending upon the structure targeted by the algorithm. We hypothesized that a nonlinear manifold learning algorithm designed to handle noisy, high-dimensional biological data, would be best suited for the signal and intrinsic noise of fMRI data. Potential of heat diffusion for affinity-based transition embedding (PHATE)

[61] is a diffusion-based manifold learning method that models local and global structures simultaneously in nonlinear dimensions. Brain activity measured with fMRI is highly noisy in both space and time, with the BOLD signal canonically peaking 4-5 s after stimulus onset before slowly returning to baseline. With a temporally-dependent stimulus such as a movie, where conversations and plot lines play out over different timescales, the autocorrelation of nearby timepoints will likely extend beyond the curve of the BOLD signal and vary by ROI along the hierarchy of temporal integration in the brain [28]. We estimate an autocorrelation kernel for each ROI by correlating each voxel timeseries with lagged versions of itself until the correlation drops and stays below zero, then averaging the functions across voxels to get a regional autocorrelation function. This function is then expanded to calculate the transitional probability between all pairs of timepoints based solely on their estimated autocorrelation and then combined with the PHATE-based transitional probability matrix and embedded in lower dimensions (Fig. 1A).

#### 5.1.1 PHATE

Given a dataset of voxel time-series data, 𝒳 = *x*_1_, *x*_2_, …, *x*_*T*_, where *x*_*t*_ ∈ R^*n*^ at time *t* is a *n*-dimensional vector and *n* is the number of voxels. Construction of the PHATE diffusion geometry starts by computing the Euclidean distance matrix *D* between data pairs, where *D*(*i, j*) = ||*x*_*i*_− *x*_*j*_||^2^.*D* is then converted into a local affinity matrix *K* using an adaptive bandwidth Gaussian kernel. The affinity matrix *K* is then row-normalized to get the initial probabilities *P*_*D*_, which are used for the Markovian random-walk diffusion process. The diffusion operator *P*_*D*_ is then raised to the *t*_*D*_th power, where *t*_*D*_is the PHATE optimal diffusion time scale, to perform a *t*_*D*_-step random walk over the data graph. This specific diffusion process infers more global relations beyond the local affinities. To further preserve both local and manifold-intrinsic global structure of data, PHATE computes the diffusion potential *U* by taking the logarithm of 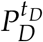and the diffusion potential distance *D*_*U*_ using distance between row pairs of *U*. This diffusion potential distance can be reduced to 2-3 dimensions for visualization or to any other dimensionality (such as that obtained by cross-validation) by performing metric MDS (MMDS).

#### 5.1.2 T-PHATE

We designed T-PHATE as a variant of PHATE that uses a dual-view diffusion operator to embed timeseries data in a low-dimensional space. The first view, *P*_*D*_, of the diffusionoperator uses the same process as PHATE to build an affinity matrix based on the Euclidean distance between data samples (here, timepoints) and then row-normalize it to obtain the transition probability matrix, where *P*_*D*_ is a *TxT* matrix. The second view *P*_*T*_ is based on an affinity matrix that summarizes the autocorrelation function of the data [62]. First we calculate a direct estimator of the autocovariance of the timeseries vector *V* from each voxel using | *V*| − 1 lags. Then the functions are averaged across voxels to obtain a single autocorrelation function (*ac f*) and smoothed with a rolling average over *w* timepoints, to account for possible jittering around where *ac f* = 0. Next, we find the first lag *lag*_*max*_ where *ac f* = 0, which defines the maximum width of smoothing for the temporal affinity matrix *A*. The temporal affinity matrix *A* is calculated as *A*(*i, j*) = *ac f* (*i, j*), 0 *<*| *i*− *j* |*>* = *lag*_*max*_ and zero elsewhere. The autocorrelative probability *P*_*t*_ is obtained by row-normalizing the affinity matrix *A*. These views are combined with alternating diffusion into a single operator *P* = *P*_*D*_*P*_*T*_, which is used as the initial probabilities for diffusion. The rest of constructing diffusion geometry and data visualization is performed the same as in PHATE. This dual-view diffusion step allows T-PHATE to learn data geometry and latent signals that represent cognitive processes that play out over longer temporal windows.

We compare T-PHATE’s performance at learning dynamic neural manifolds with common dimensionality reduction algorithms that are agnostic to time including principal components analysis (PCA), uniform manifold approximation and projection (UMAP) [38], and vanilla PHATE [61]. To test whether our autocorrelation kernel was the best approach to modeling temporal dynamics in this manifold, we tested two additional versions of PHATE: incorporating time as a feature in the input data (PHATE+Time) and smoothing the data temporally (Smooth PHATE). fMRI data were extracted from regions of interest (ROIs) and normalized before embedding.

### 5.2 Event segmentation modeling

Human experience of real life is continuous, yet perception and conception of this experience is typically divided into discrete moments or events. Event segmentation theory explains that humans automatically divide continuous streams of information into discrete events [45, 47] in order to form, organize, and recollect memories, make decisions, and predict the future [63]. Participants show high consistency in explicitly segmenting continuous stimuli [46] and also in how their brains represent these event boundaries [50, 58]. Event boundaries can be represented, both behaviorally and in brain activity, at different timescales depending on the information being used to draw event boundaries [57]. In the brain, event boundaries are reflected by shifts in the stability of regional activity patterns, and we can learn the structure of events with a variant of a Hidden Markov Model [50]. During a continuous stimulus such as a movie, different brain regions represent events along different timescales, which reflect the dynamics of the information being represented by a given region. For example, early sensory regions represent low-level information about a stimulus. In most movie stimuli, low-level sensory features shift quickly as cameras change angles or characters speak, and so early sensory regions show more frequent event shifts [50, 52]. Later sensory regions represent longer timescale events, such as conversations and physical environments that change less frequently than features like camera angles. Regions associated with memory or narrative processing represent events on longer timescales, and these boundaries best correspond with scene changes marked by human raters [51].

We used the Hidden Markov Model (HMM) variant presented in [50] and implemented in BrainIAK [64] to learn from different representations of BOLD timeseries data where a brain region experiences event boundaries, or transitions between stable states of activation. Given an activation timeseries of a brain region during a continuous stimulus, the HMM identifies stable activity patterns or “events” that are divided by boundaries, where the timeseries transitions between two stable patterns of activation. This is done iteratively and the log-likelihood of the model fit is evaluated at each iteration, and model fitting stops when the log-likelihood begins to decrease. The first step of this process is to use the HMM to learn from the data the number of events a brain region represents for a given stimulus. Past studies have used HMMs for event segmentation on multivoxel activity patterns and have validated this approach against behavioral segmentation [50, 51, 58]. This shows that voxel resolution data reflect meaningful event segmentation, so we chose to estimate the optimal number of events (*K*) for each brain region using the voxel resolution data (Fig. 4), which also prevents overfitting to the embedding data.

To run an HMM for event segmentation on our manifold embedding data, we needed to tune two parameters for each subject: *K*, or the number of hidden states through which a brain region passes during the stimulus, and *M*, the dimensionality of the latent space that captures this signal.

#### 5.2.1 Optimizing number of neural events

We estimated *K* and *M* for each brain region and embedding method with a leave-one-subject-out cross-validation scheme. Using BOLD data from *N*− 1 training subjects, we created an average timeseries matrix by averaging response profiles for each voxel across the training subjects. We define *K* as the number of hidden states in an HMM and optimize *K* by fitting a model with *K* states on the training data, applying it on the BOLD data of the held-out test subject, and calculating the log probability of the test data according to the learned event segmentation of the model (hence referred to as the model log-likelihood). We tested a range of *K* from 2 to 200 for *Sherlock* and 2 to 400 for *Study-Forrest*. For each training fold, we calculated the best *K* value that maximized the model log-likelihood of the voxel resolution data, setting subject *N*’s *K* value to the average of the *N*− 1 training subjects’ *K*s. After definition on the voxel resolution data, *K* for each subject and region was held constant for all embedding types.

#### 5.2.2 Optimizing manifold dimensionality

We used the same leave-one-subject-out cross-validation scheme to select the dimensionality of manifold embeddings for each subject. We embedded each subject’s timeseries data into an array of possible *M* values then fit an HMM on each *M*-dimensional embedding with the subject’s cross-validated *K* value, which had been learned in the previous step from all the other subjects’ voxel resolution data. Event boundary fit evaluation is described in more detail below. The *M* parameter that maximizes the event boundary fit was recorded for each test subject. We then set the *M* value to be used for subject *N* as the average of all *N* − 1 subjects’ best *M* values.

#### 5.2.3 Evaluating the fit of event boundaries

To quantify how well a learned manifold embedding amplifies dynamic structure in the data, we calculated difference in correlation across latent dimensions for pairs of time-points within and across event boundaries. For this calculation, we restricted the time-points that went into the calculation as follows. We calculated the length of the longest event, and only considered timepoints that were less than that temporal distance apart. We anchored a comparison on each timepoint *t* in the timeseries, and for each temporal distance *n* ranging from 1 timepoint to the length of the longest event, we considered a pair of timepoints *t*− *n* and *t* + *n* if one timepoint fell in the same event as the anchor and the other a different event. We then took the correlation between the anchor and each timepoint, binning them according to whether they were a within-event comparison or a between-event comparison. This process assures that there are equal numbers of timepoints going into each comparison type (Fig. 5A).

One issue of defining and testing event boundaries within-subject is overfitting. To circumvent this, we cross-validated the event boundaries across subjects, testing *N*−1 sets of event boundaries on each *N*^*th*^ subject’s neural data using the same procedure outlined above.

To assess the behavioral relevance of the neural event segmentation, we used a set of event boundaries identified by a separate cohort of *Sherlock* study participants. These participants watched the *Sherlock* episode outside of the scanner and were asked to indicate where they believed a new event began [65] (Fig. 7). We applied these human-labeled boundaries to the neural data and measured the fit of the boundaries (as outlined above) to gauge how the embeddings not only highlight neural dynamics but how those neural dynamics relate to the conscious, real-world experience of events.

### 5.3 Data

#### 5.3.1 Sherlock dataset

For full details on the *Sherlock* data, see the original publication of this dataset [54]. Here, we used data from the 16 participants who viewed a 48-minute clip of the BBC television series “Sherlock.” Data were collected in two fMRI runs of 946 and 1030 time-points (repetition time; TRs) and was downloaded from the DataSpace public repository (http://arks.princeton.edu/ark:/88435/dsp01nz8062179). The data were collected on a Siemens Skyra 3-T scanner with a 20-channel head coil. Functional images were acquired with a T2*-weighted echo-planar imaging sequence (TE 28ms, TR 1.5s, 64-degree flip angle, whole brain coverage with 27 slices of 4mm thickness and 3×3 mm^2^ in-plane resolution, 192×192mm^2^ FOV). Anatomical images were acquired with a T1-weighted MPRAGE pulse sequence with 0.89mm^3^ resolution. Slice-time correction, motion correction, linear detrending, high-pass filtering (140s cutoff), and co-registration and affine transformation of functional volumes to the Montreal Neurological Institute (MNI) template were all performed with FSL. Functional images were then resampled from native resolution to 3mm isotropic voxels for all analyses, z-scored across time at every voxel, and smoothed with a 6mm kernel.

#### 5.3.2 StudyForrest dataset

For full details on the *StudyForrest* data, see the original publication of the movie dataset [66] and the localizer extension dataset [67]. Here, we included data from 15 participants who completed both tasks. All participants were native German speakers. In the movie-viewing task, participants watched a 2-hour version of the movie “Forrest Gump.” These data were collected in eight fMRI runs resulting in a full timeseries of 3599 TRs (451, 441, 438, 488, 462, 439, 542, and 338 per run). Movie data were collected on a 7-T Siemens scanner with a 32-channel headcoil. Functional images were acquired with a T2*-weighted echo-planar imaging sequence (TE 22ms, TR 2s, whole brain coverage with 36 slices of 1.4 mm thickness and 1.4 × 1.4 mm^2^ in-plane resolution, 224mm FOV). In the localizer task, the same fifteen participants viewed 24 unique grayscale images from 6 categories (human faces, human bodies without heads, houses, small objects, outdoor scenes, and scrambled images) in four (156 TR) block-design runs with two 16s blocks per stimulus category per run. Localizer data were collected on a 3-T Philips Achieva scanner with a 32-channel headcoil. Functional images for the localizer task were acquired with a T2*-weighted echo-planar imaging sequence (TE 30ms, TR 2s, 90-degree flip angle, 35 slices of 3mm thickness and 3×3 mm^2^ in-plane resolution, 240mm FOV). Structural images were acquired with a 3-T Philips Achieva using a 32-channel headcoil. T1-weighted anatomical images were acquired with a 3D turbo field echo sequence with 0.67mm isotropic resolution. Slice-time correction, co-registration and affine transformation to MNI template were performed with FSL where functional images were resampled to 3mm isotropic voxels. Additional preprocessing included linear detrending, high-pass filtering (100s cutoff), spatial smoothing with a 6mm kernel, and nuisance regression (including 6 motion parameters, global signal, white matter, and cerebrospinal fluid), and z-scoring within voxel to match the *Sherlock* preprocessing.

### 5.4 Region of interest (ROI) selection

We selected four regions of interest (ROIs) based on the original publication of the *Sherlock* dataset [54], subsequent publications [50], and regions known to have reliably strong signals in response to audiovisual movie stimuli [68]. The early visual (EV), early auditory (EA), and high visual (HV) region masks were based on a functional atlas defined with resting-state connectivity [69]. As in the original *Sherlock* publication, we defined a posterior medial cortex (PMC) ROI as the posterior medial cluster of voxels within the dorsal default mode network [54]. In the *Sherlock* data, the dimensionality of voxel-resolution ROIs are as follows: EV = 307, HV = 571, EA = 1018, PMC = 481. In the *StudyForrest* data, the dimensionality of voxel-resolution ROIs are as follows: EV = 166, HV = 456, EA = 657, PMC = 309.

## 5.5 Code availability

T-PHATE is available as a python package at: https://github.com/KrishnaswamyLab/TPHATE. The pipeline to replicate all analyses presented here is available at: https://github.com/ericabusch/tphate_analysis_capsule.

## 5.6 Data availability

The *Sherlock* dataset was downloaded from the Dataspace Public Repository at the following link: http://arks.princeton.edu/ark:/88435/dsp01nz8062179. The *StudyForrest*dataset was accessed via DataLad [70] from here: https://github.com/psychoinformatics-de/studyforrest-data. Steps to reproduce our preprocessing pipeline and ROI extraction are available here: https://github.com/ericabusch/tphate_analysis_capsule.

## 5.7 Author contributions

E.L.B., J.H., G.W., G.L., S.K., N.B.T-B. conceived the idea. E.L.B., J.H., A.B., and S.K. designed the T-PHATE algorithm. E.L.B. and J.H. developed software. E.L.B. curated data and designed analyses, and E.L.B, J.H., and T.W. performed the analyses. E.L.B., S.K., and N.B.T-B. wrote the initial manuscript draft. All authors revised the paper. S.K. and N.B.T-B. jointly supervised the work.

## 1 Supplemental methods

### 1.1 Movie annotations

#### *Sherlock* annotations

We used annotations for the *Sherlock* movie stimulus released with [65] and available freely on GitHub: https://github.com/kiranvodrahalli/fMRI_Text_maps_NI. These annotations were created by human annotators who viewed the film and labeled each timepoint on a variety of metrics. Specifically here we used two sets of binary labels that correspond with visual and auditory aspects of the stimulus: whether a given TR featured an indoor or outdoor scene and whether or not the TR featured music.

#### *StudyForrest* annotations

Annotations of scene cuts and locations used in later analysis of the Forrest Gump movie viewing data were downloaded from a subsequent *StudyForrest* publication [71]. These provide additional information about a number of shots from the film, including the setting of the shot, whether a setting is interior or exterior, the temporal progression of a shot with reference to the previous shot, and the time of day. These are used in the movie feature decoding analysis for the *StudyForrest* dataset.

## 2 Supplemental results

**Figure S1:**
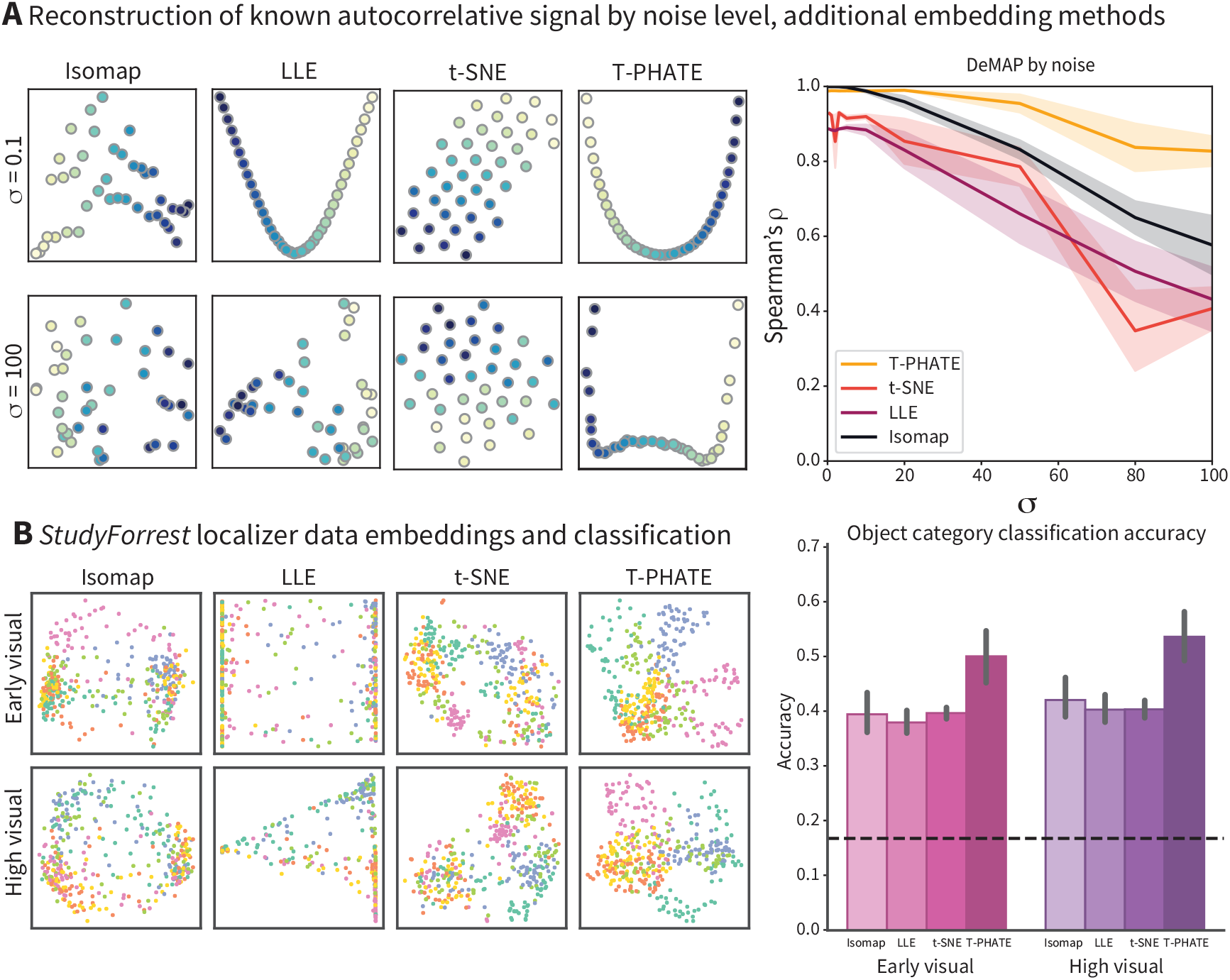
**A**. As in figure 1B, we simulated data with a known autocorrelative function and varied the error added to the signal via normally-distributed noise with varying *σ*. We replicate the results of 1B on three additional benchmarks: isometric mapping (isomap), locally linear embeddings (LLE), and t-distributed stochastic neighbor embedding (t-SNE). **B**. We visualized 2-dimensional embeddings of the *StudyForrest* localizer data for a sample subject (S2) and performed support vector classification to predict object category from these embeddings, as in figure 2.

**Figure S2:**
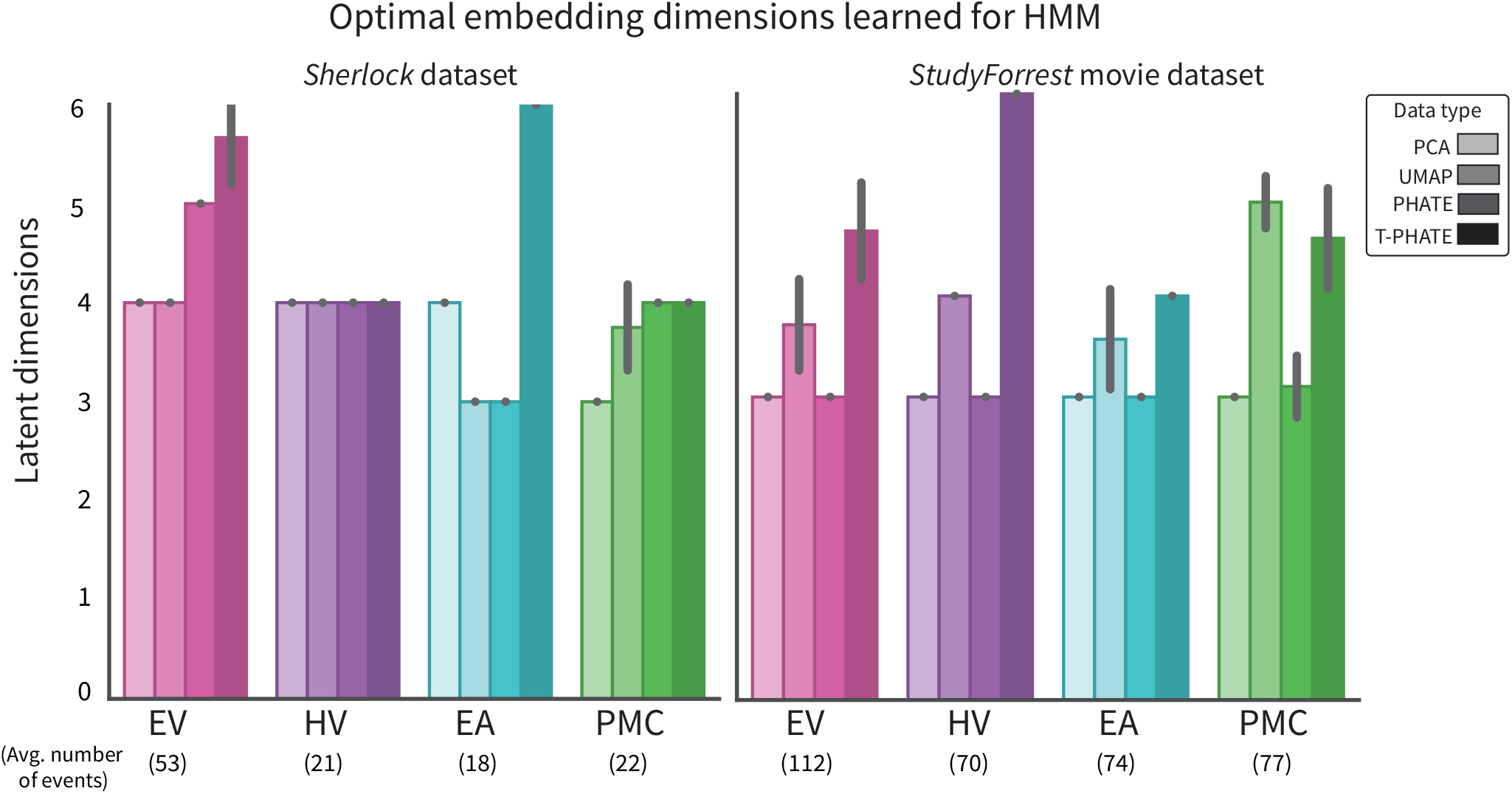
HMM parameters *K* and *M* learned with leave-one-subject-out cross validation. Dimensionality reduction methods are able to optimize event boundaries in relatively few dimensions. *K* values follow the predicted event hierarchy in the brain, with lower-level sensory regions experiencing more events than higher-order cognitive regions.

**Figure S3:**
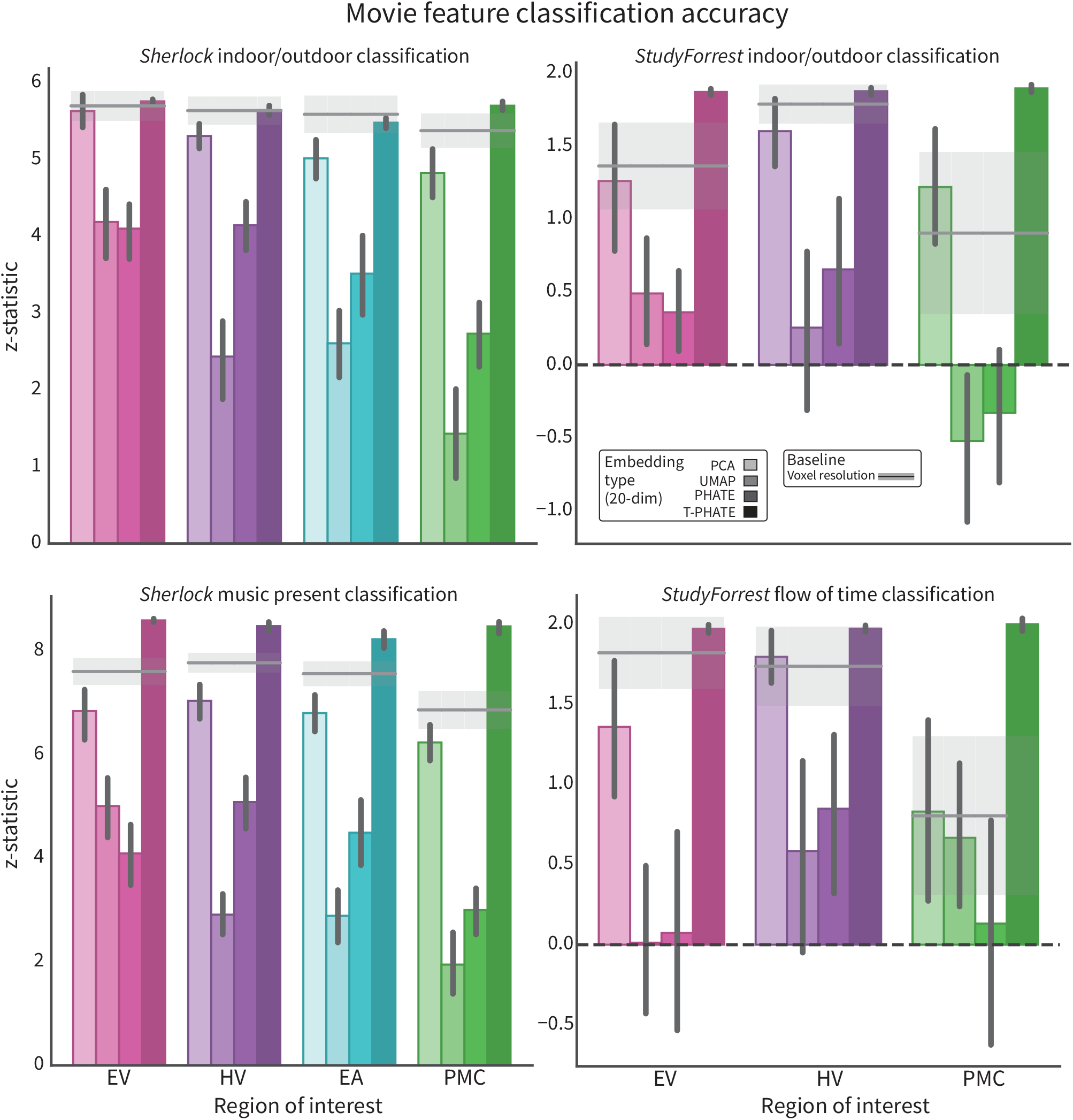
Support vector classification of movie features in the *Sherlock* and *StudyForrest* movie datasets. All labels were binary aside from the Flow of Time labels, which had four levels. Results presented are the z-statistic of true classification accuracy versus 1000 repetitions of time-shifting labels and re-training the model to generate a null distribution.

**Figure S4:**
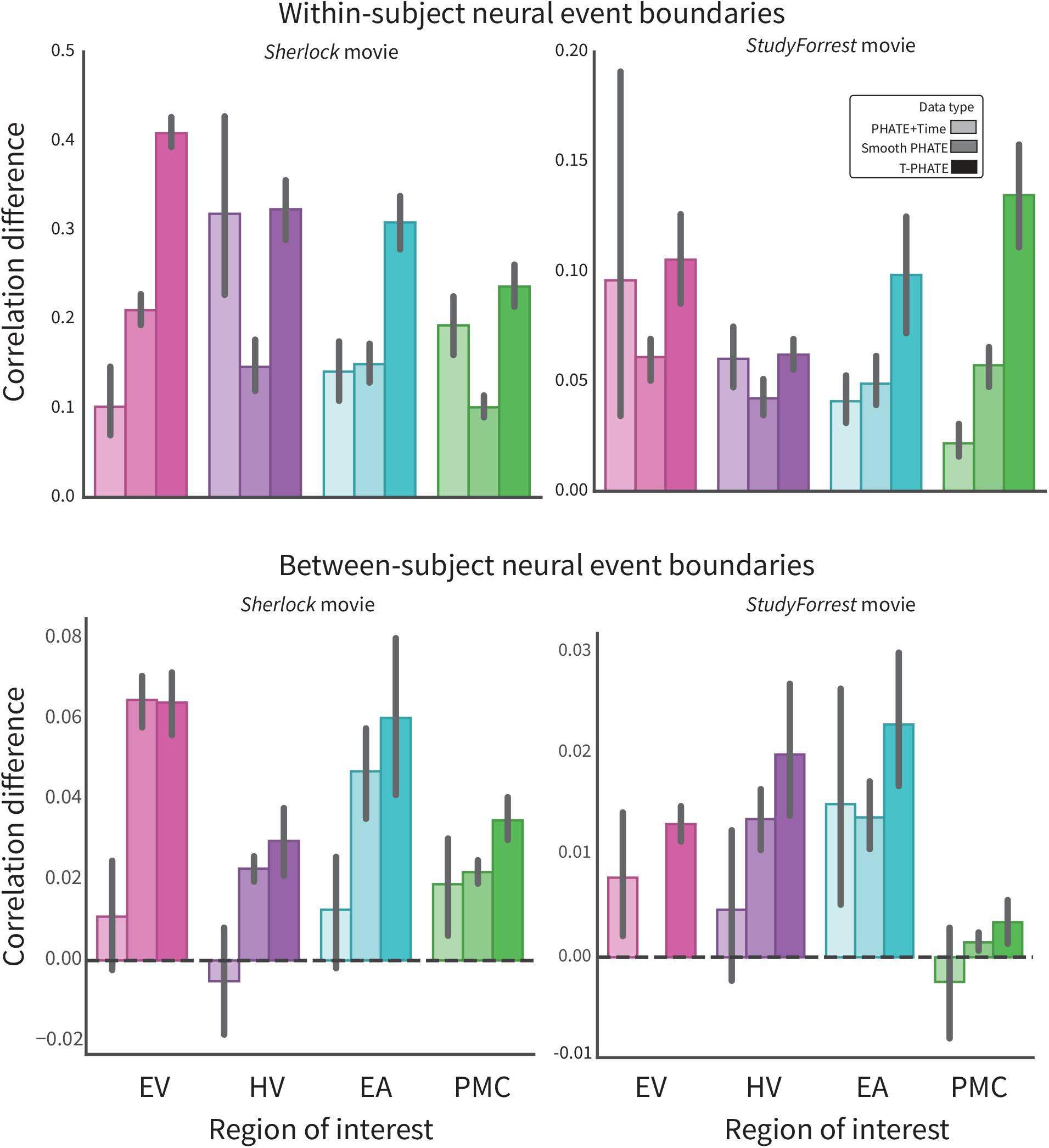
Within-vs-between neural event boundaries for the time alternative methods. **A**. Event boundaries defined as neural events within-subject. **B**. Event boundaries cross-validated across subjects with leave-one-subject-out boundaries.

**Figure S5:**
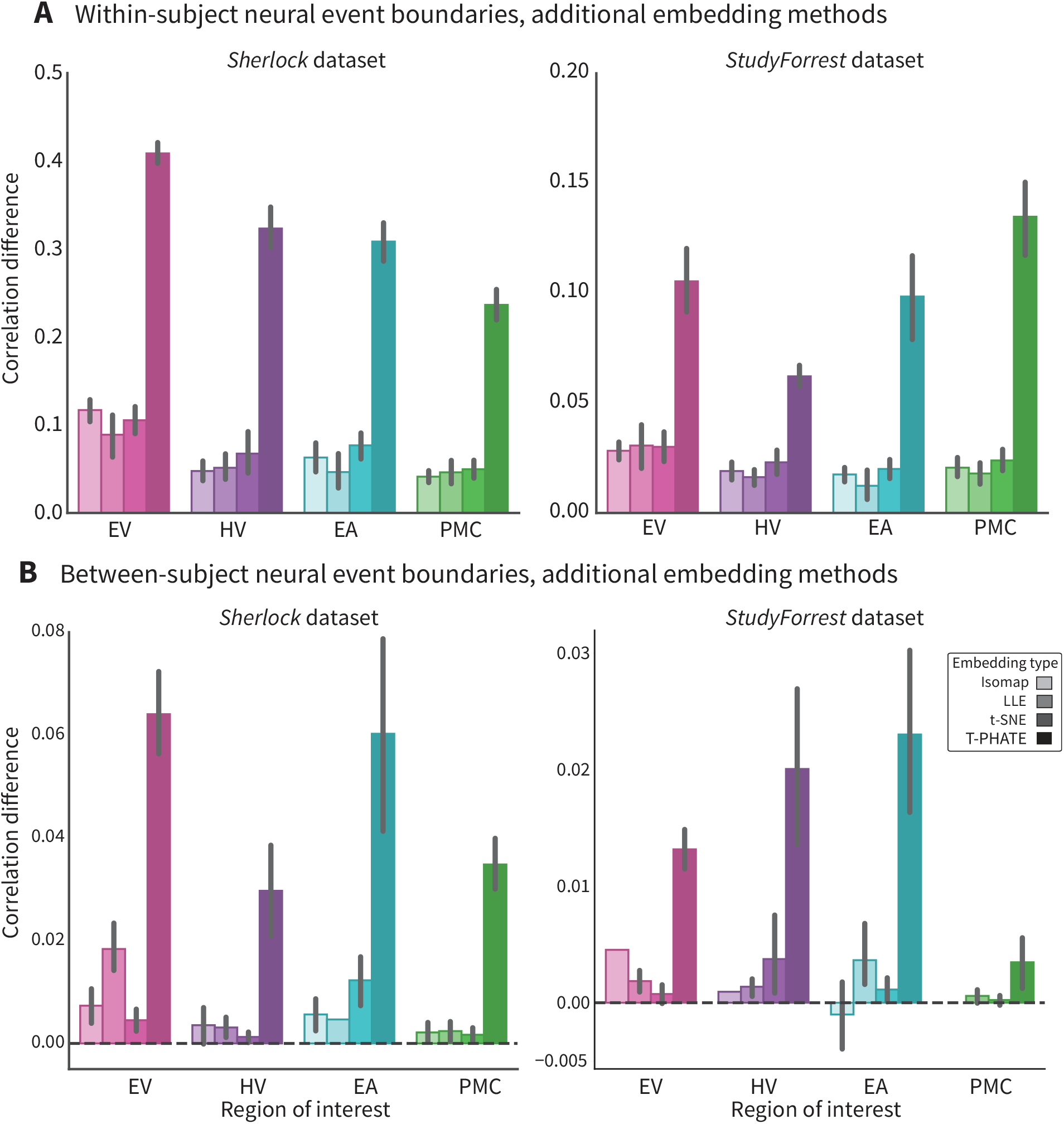
Within-vs-between neural event boundaries for Isomap, LLE, and t-SNE. **A**. Event boundaries defined as neural events within-subject. **B**. Event boundaries cross-validated across subjects with leave-one-subject-out boundaries.

